# Municipal Wastewater in Ottawa, Canada contains a reservoir of bacteriophages with clinically relevant *Escherichia coli* and *Klebsiella pneumoniae* hosts

**DOI:** 10.64898/2026.09.20.752992

**Authors:** Sean Stephenson, Walaa Eid, Elizabeth Mercier, Robert Delatolla, Adam D. Rudner, Tyson E. Graber

**Author notes:** Correspondence: Tyson E. Graber. **Repositories** The newly sequenced viral genomes have been deposited in the GenBank database under accession numbers PZ683200, PZ683201, PZ683202, PZ683203, PZ683204, PZ683205, PZ683206, PZ683208, PZ683209, PZ683207.

## Abstract

The recent rise of antimicrobial resistant bacteria in health care and environmental settings has been detrimental to the feasibility of antibiotic treatments for patients and agricultural applications. Along with this escalating issue, a resurgence of bacteriophage research has occurred, and phage therapy is proving to be a powerful case-specific alternative. However, a centralized source of phages that are accessible, abundant, and characterized as candidates for potential use in therapeutic applications remains to be fully established. This study aims to identify the range of bacteriophages available in municipal wastewater from Ottawa, Ontario that can infect and lyse *Escherichia coli* O1:K1:H7 and *Klebsiella pneumoniae* K3 hosts. Building-level sewer shed, influent, and primary sludge samples were screened for the presence of these phages which were then isolated through several rounds of plaque purification, sequenced by Nanopore technology, and annotated. Ten phages belonging to the genus *Kayfunavirus, Vectrevirus, Kuravirus*, or *Sugarlandvirus* were identified and bio-banked from different types of wastewater at several time points. These phages are 96-99% identical to phages isolated in Germany, Switzerland, and Spain, but contain divergent structural and metabolic genes. This work contributes to the ever-growing database of sequenced phages, provided with a physical archive of the organisms, and helps uncover the extent of phage biodiversity at different levels of wastewater treatment facilities that are readily available for screening in phage therapy applications.

## 3. Introduction

A rebirth of bacteriophage therapy has taken place over the last quarter century due to the rise of multidrug-resistant strains of bacteria that greatly reduce the general efficacy of antibiotics (1,2). These viruses selectively infect bacteria causing the arrest and lysis of cultures in vitro (3) and an increasing amount of evidence is showing very promising results when using them to treat human illnesses. Bacterial infections of the urinary tract, lungs, and those involving surgically implanted devices are among the most common conditions that have been treated with bacteriophage therapy to date (4,5). Many applications outside of human health, such as agriculture and food safety, are also gaining recognition, but local and national repositories of accessible bacteriophages still need to be developed to expedite the full potential of phage therapy (6).

Extraintestinal pathogenic *E. coli* strains cause a wide range of illnesses due to infections occurring outside of the gut. This classification of *E. coli* is based on the symptoms of infected patients, multilocus sequence typing, and serotype (7). Molecular serotyping is used for the pathogenic analysis of *E. coli* by assigning strains into groups based on the sequences corresponding to three main antigens; the lipopolysaccharide membrane (O), capsule (K), and flagella (H) (8,9). *E. coli* O1:K1:H7 is a clinically relevant serotype that causes urinary tract infections, sepsis, and neonatal complications all over the world (10–12). In addition to its problematic nature in humans, strains with the same serotype and homologous genomes have been characterized as avian pathogenic *E. coli*. These strains have been found to infect the lungs of chickens creating a potential link between animal and humans through poultry (13,14). Closely related O2:K1, in addition to O1:K1 strains, have also been shown to infect neonatal rats and highlight the zoonotic potential of this pathogen (15). Compounding all these effects is the rise of antibiotic-resistant forms of *E. coli*, particularly in urinary tract infections. The presence of beta-lactamase and mobile colistin resistance genes have been on the rise in sequenced clinical isolates over the last decade (16,17). Although not explicitly found in serotype O1:K1:H7, the acquisition of these or similar genetic elements is a concern due to the plasmid-borne nature of these resistance genes and horizontal gene transfer being a common occurrence in *E. coli*. (18).

Hypervirulent strains of *K. pneumoniae* are responsible for various hospital-acquired infections worldwide, such as liver abscess, meningitis, and pneumonia (19,20). There are many different strains of *K. pneumoniae* that are mainly categorized by their capsule (K) type. Over 70 serotypes exist based on antigens related to these outer membrane proteins, but only a subset is responsible for pathogenesis in humans (21,22). Serotypes K1, K2, K5, K20, K54, and K57 are among the most virulent, however a combination of other biomarkers including antibiotic resistance genes are being increasingly used for classification and diagnosis (23,24). Antibiotic resistance is growing hallmark of virulent *K. pneumoniae* strains and is causing a wide panel of drugs to lose efficacy (25,26). A large driver of strain diversity is horizontal gene transfer through plasmids that contain antibiotic resistance genes, such as *bla_KPC_* and beta-lactamase (27–30).

This study describes a snapshot of the landscape of *E. coli* O1:K1:H7 and *K. pneumoniae* K3 bacteriophages that can be found in municipal wastewater sampled in from Ottawa, Canada, a city of 1 million inhabitants. An assessment of the diversity of species based on their sequence and morphology was conducted to explore the feasibility of using primary sludge, influent, or building-level wastewater to easily isolate a broad range of bacteriophages that have potential uses in clinical and agricultural applications.

## 4. Methods

### Cultivation of bacterial strains

Bacterial strains *Escherichia coli* O1:K1:H7 (ATCC11775) and *Klebsiella pneumoniae* K3 (ATCC13883) were cultured in Nutrient Broth (Millipore #70122) with 0%, 7.5%, or 15% agar (BD #214010) for liquid, overlay, and solid media respectively. Cultures were grown at 37°C overnight, and with shaking at 120 rpm for liquid media. All procedures were performed in Containment Level 2 and approved by the CHEORI Biosafety Committee.

### Isolation of bacteriophages from wastewater

Primary clarifier sludge and influent (post grit treatment wastewater samples) were obtained from the Robert O. Pickard Environmental Centre, the main wastewater resource recovery facility in Ottawa, Canada, and stored at 4°C until processing. Primary sludge samples were diluted 1/20 in phage buffer (10 mM Tris-HCl pH 7.5, 10 mM MgSO_4_, 68 mM NaCl) then treated as influent throughout the rest of the preparation. A single 24-hour composite wastewater sample was taken from a residence building at the University of Ottawa and treated as influent. Influent and diluted primary sludge samples were centrifuged for 10 minutes at 3,000 rcf to pellet bacteria and debris, and the resulting supernatant was sequentially passed through a 0.45 µm and 0.22 µm pore sized PES membrane filter. Filtrate was then used in downstream plaque assays or stored at 4°C.

### Isolation and purification of bacteriophage stocks

Filtrates from wastewater were mixed with overnight bacterial culture and melted agar media warmed to 55°C before being plated using standard top-overlay technique and incubated at 37°C overnight. Plaques were picked using a sterile pipette tip and resuspended in phage buffer. Serial dilutions of resuspended phage were plated again to obtain isolated plaques, and then this process was repeated twice more. After purification, high density (webbed plates) were prepared, flooded with phage buffer, and incubated at room temperature for 4 hours. Phages were harvested by removing the phage buffer from the plate, centrifuging at 3,000 rcf for 20 minutes, and passing the supernatant through a 0.22 µm filter. The resulting filtrate was stored at 4°C until further use.

### DNA extraction

DNA for restriction digest and sequencing was extracted from high titre (>10^9^ pfu/mL) bacteriophage stocks using PCI (25:24:1 phenol:chloroform:isoamyl alcohol). Prior to extraction, bacteriophage stocks were supplemented with MgCl_2_ (2.5 mM) and CaCl (0.5 mM) then treated with DNase I (2 U/mL) and RNase A (10 µg/mL) for 1 hour at 37°C. Following nuclease treatment, Tris-HCl (10 mM, pH 7.5), PEG (10%) and NaCl (200 mM) were thoroughly mixed into the solution then incubated at 4°C overnight. After precipitation, viral particles were pelleted at 12,000 rcf for 20 minutes at 4°C and then resuspended in phage buffer. Resuspended bacteriophages were treated with proteinase K (50 µg/mL) in the presence of 0.5% SDS for 1 hour at 55°C with intermittent mixing to digest capsids. Next, an equal volume of PCI was added and mixed vigorously by shaking before separating the phases by centrifugation at 16,000 rcf. The aqueous top phase containing DNA was mixed with an equal volume of chloroform, and the process used for PCI was repeated. The second aqueous phase was mixed with 2 volumes of ice-cold ethanol (100%) and NaOAc (100 mM) then stored at -20°C overnight to precipitate DNA. The DNA was pelleted by centrifugation at 16,000 rcf for 10 minutes at 4°C, and the pellet was washed with 70% ethanol. After another round of centrifugation, the ethanol was removed and the pellet was air-dried for 15 minutes in a fume hood then finally resuspended in pure water. Genomic DNA from bacteriophages was digested using either AluI, ClaI, DraI, NotI, PacI, or XhoI (New England Biolabs) in 1X CutSmart buffer for 30 minutes at 37°C. Uncut DNA and digestion reactions were loaded onto a 0.7% agarose gel made in 1X TAE with 1X GelRed (Biotium) using 6X loading dye. Gels were run for 45 minutes at 100 V then imaged using a Bio-Rad Gel Doc.

### Genome sequencing and bioinformatics

Complete genomes were obtained using Oxford Nanopore sequencing. Direct library preparation was achieved using the Rapid Barcoding Kit 24 V14 (SQK-RBK114.24) according to manufacturer instructions and sequences were read using the MinION Mk1B with a FLO-MIN114 Flow Cell. Base calling and read quality were assessed by MinKNOW (v25.03.9) and further evaluated by NanoPlot (v1.46.2) (31) before assembly with Flye (v2.9.6) (32) and polishing of consensus sequences with medaka (https://github.com/nanoporetech/medaka). Annotation was conducted using Pharokka (v1.8.2) (33) followed by Phold (v1.2.2) (34), and genus classification was determined with ViPTree (v4.0) (35) and BLASTn (36) against the core nucleotide database. The location of terminal repeats was validated using existing reports of closely related phages.

### Transmission Electron Microscopy

Images of phages were obtained through transmission electron microscopy. Tween-20 was added to phage samples at 0.01% and incubated on carbon-coated copper grids (Ted Pella, #01813) for approximately 5 minutes, before being removed by wicking. Grids were washed twice with 0.01% Tween-20 and stained with 1% uranyl acetate for 2 minutes. Stained grids containing bacteriophages were imaged using the JEOL JEM-1400Flash (80 kV) under vacuum at 80kX magnification. Scale bars were added and measurements were made using ImageJ software.

## 5. Results

### Bacteriophages isolated from urban wastewater in Ottawa

Ten bacteriophages (CROW1-10) were independently isolated at the <u>C</u>HEO <u>R</u>esearch Institute from the city of <u>O</u>ttawa’s <u>w</u>astewater located within Ontario, Canada. Three distinct types of wastewater from two separate locations were screened for the presence of bacteriophages: influent and primary sludge sampled from the city’s main water resource recovery facility (Robert O. Pickard Environmental Centre, ROPEC), and building-level wastewater sampled in the University of Ottawa sewer shed. A total of seven bacteriophages that infect *Escherichia coli*, and three bacteriophages with *Klebsiella pneumoniae* as a host were identified (Table 1). All bacteriophages were isolated directly using small volumes of each sample without the need for enrichment with host bacteria.

**Table 1.**
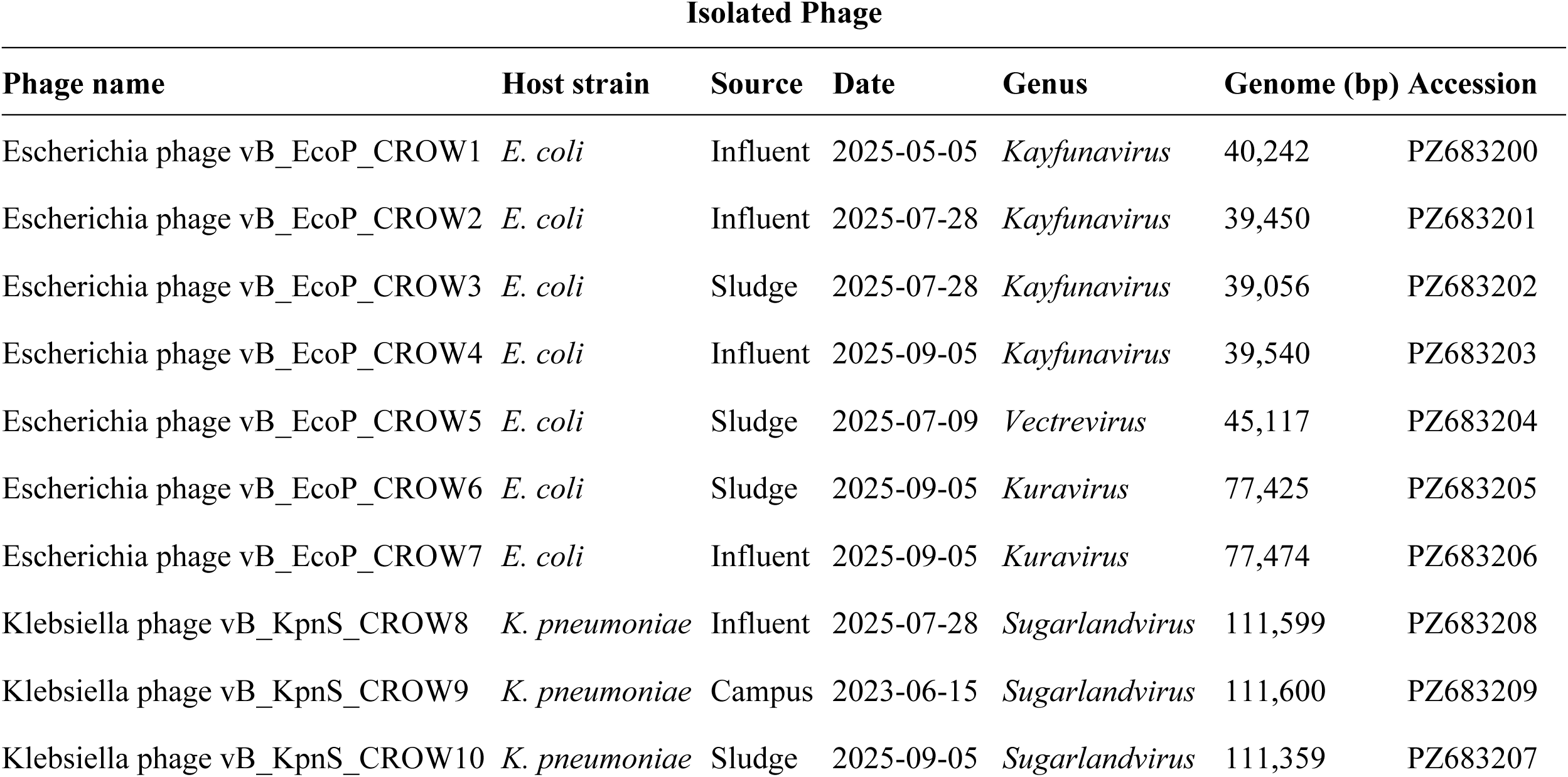
Isolated phages.

### Bacteriophage and plaque morphology

After three rounds of purification, the plaque morphology produced by each bacteriophage population was observed. Plaques were variable in size and appearance, showing a mix of turbidity and lysis patterns on top-agar overlays (Figure 1, left panels). All bacteriophages isolated with the *Escherichia coli* O1:K1:H7 host generated plaques with bullseye lysis patterns and ranged in size from approximately 2-5 millimeters in diameter. Plaques produced from bacteriophages with the *Klebsiella pneumoniae* host were notably smaller (approximately 1 millimeter diameter or less) and had no observable lysis patterns.

**Figure 1.**
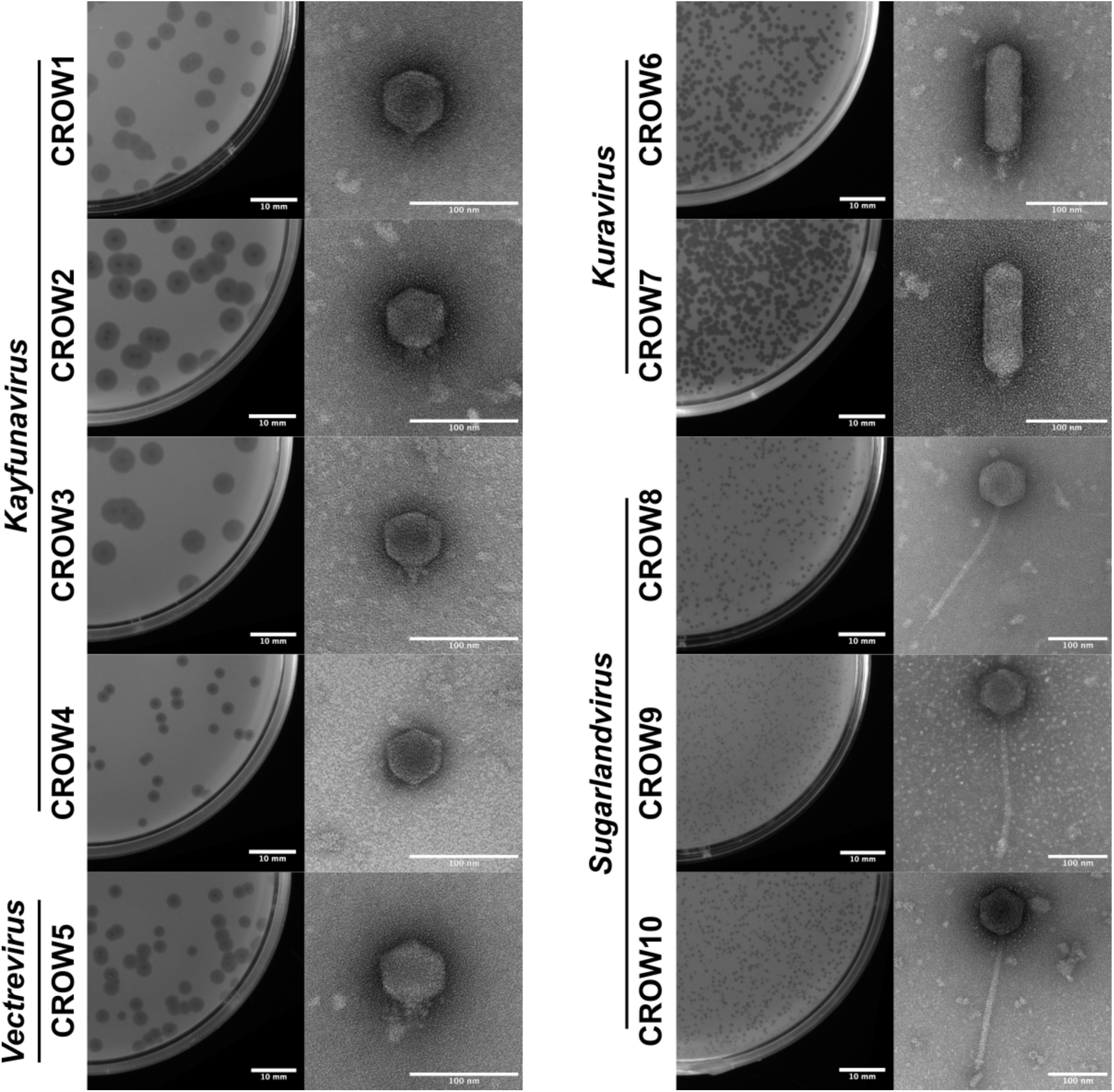
Bacteriophage morphology. Ten bacteriophages are shown as plaques (left panels) and viral particles (right panels). Plaques appearing on bacterial lawns of *E. coli* and *K. pneumoniae* were visualized under white light without magnification (scale bar = 10 mm). Viral particles were negatively stained with uranyl acetate and imaged using transmission electron microscopy at 60,000-80,000x magnification with an accelerating voltage of 80kV (scale bar = 100 nm).

Each bacteriophage was propagated, and high titre stocks were subject to transmission electron microscopy for morphological analysis. Imaging of viral particles revealed some well-studied and other more unique morphologies belonging to either the *Siphoviridae* or *Podoviridae* family (Figure 1, right panels). Five of these phages (CROW1, CROW2, CROW4, CROW3, CROW5) shared the most common morphology for this family with small icosahedral heads approximately 60 nm in diameter and short tail fibers. The other two *Podoviridae* (CROW6 and CROW7) displayed a rarer morphotype with prolate heads approximately 130 nm in length and 50 nm in width, with similarly short tail fibers. All three phages isolated with the *Klebsiella pneumoniae* host (CROW8, CROW9, and CROW10) appeared to have classical *Siphoviridae* morphology with an icosahedral head approximately 80 nm in diameter and a tail roughly 250 nm long. Interestingly, for bacteriophages isolated with the *Escherichia coli* host, two distinct *Podoviridae* morphologies were observed.

### Taxonomy analysis & novelty assessment

Further characterization of these bacteriophages was achieved through whole genome sequencing using long-read Nanopore technology. Complete genomes were assembled into a single contig, and coding sequences were then subject to proteomic analysis using ViPTree. This analysis showed that all ten phages belong to the *Pseudomonadota* host group (Figure 2), with half of the bacteriophages belonging to the order *Autographivirales*, and the rest to less established families of the *Caudoviricetes* class. Each cluster revealed several closely related phages that allowed for further comparisons and determination of each individual phage taxonomy. A complimentary analysis at the nucleotide level using BLASTn confirmed that the independently isolated bacteriophages separated into four unique clusters according to their genus. Four were classified as *Kayfunavirus* (CROW1, CROW2, CROW3, CROW4), one as *Vectrevirus* (CROW5), two as *Kuravirus* (CROW6, CROW7), and three as *Sugarlandvirus* (CROW8, CROW9, CROW10). The percent identity of these sequences was greater than 95% when compared to their closest matches in the core nucleotide BLAST database (Table 2) indicating these phages were part of an existing genus with uniqueness at the species level.

**Figure 2.**
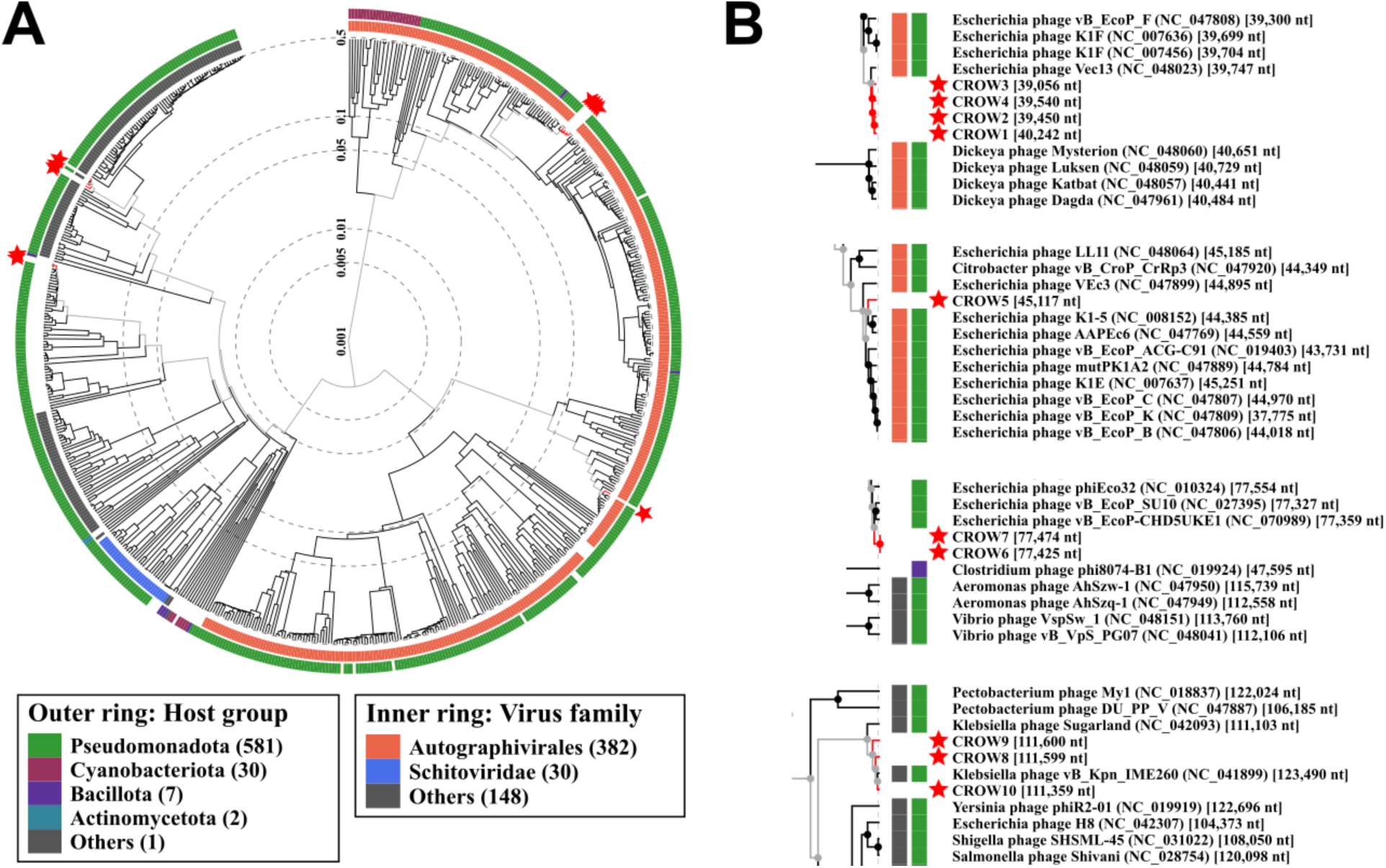
Viral proteomic tree of isolated and reference bacteriophages. (A) Virus families (inner ring) are clustered by genome-wide similarities based on translated nucleotide BLAST scores. Host groups (outer ring) are displayed for each virus along with the genomic distance scale (dashed lines, 0.001-0.5). Isolated bacteriophages are indicated with a star. (B) An expanded view of isolated bacteriophage clusters.

**Table 2.** Comparison of phage genomes using BLASTn.

| Isolated phage<br>*Kayfunavirus* | Comparison phage |  |  |  |
| --- | --- | --- | --- | --- |
|  | EcSW4 |  | K1F |  |
|  | Identity (%) | Query cover (%) | Identity (%) | Query cover (%) |
| CROW1 | 96.49% | 96% | 94.87% | 85% |
| CROW2 | 96.54% | 96% | 94.90% | 88% |
| CROW3 | 96.70% | 95% | 94.89% | 91% |
| CROW4 | 96.13% | 95% | 94.64% | 87% |
| *Kuravirus* |  |  |  |  |
|  | CHD5UKE1 |  | SU10 |  |
|  | Identity (%) | Query cover (%) | Identity (%) | Query cover (%) |
| CROW6 | 99.04% | 95% | 96.23% | 97% |
| CROW7 | 99.06% | 94% | 96.24% | 96% |
| *Vectrevirus* |  |  |  |  |
|  | K1-5 |  | K1E |  |
|  | Identity (%) | Query cover (%) | Identity (%) | Query cover (%) |
| CROW5 | 95.32% | 90% | 94.63% | 89% |
| *Sugarlandvirus* |  |  |  |  |
|  | IME260 |  | K30lambda2.2 |  |
|  | Identity (%) | Query cover (%) | Identity (%) | Query cover (%) |
| CROW8 | 97.78% | 95% | 96.72% | 98% |
| CROW9 | 96.74% | 91% | 97.30% | 92% |
| CROW10 | 96.57% | 97% | 92.80% | 96% |

### Genomic analysis of Kayfunavirus bacteriophages

The genome size of the *Kayfunavirus* bacteriophages CROW1, CROW2, CROW3, CROW4 are 40,242, 39,450, 39,056, and 39,450 base pairs respectively. Using CROW1 as an example to describe the landscape of these *Kayfunavirus* genomes, 62 ORFs were identified with 29 having an unknown function (Figure 3A). Genes corresponding to the head, tail, connector, DNA/RNA metabolism, host takeover, and lysis are present, but no tRNA genes were found. The GC content is 49.94% with periodic shifts in GC skew. The four *Kayfunavirus* phages isolated in this study had 96.6-99.6% nucleotide percent identity to one another. Belonging to an established genus of bacteriophages allowed for comparisons to what has already been reported, such as bacteriophage K1F that has an endosialidase domain within its tail spike structure (37,38). These enzymatic receptor binding proteins mediate the O-antigen expression dependant infection of hosts and have co-evolved broadly; even across morphotypes, as demonstrated by homologous pectin lyase-like tail spikes possessed by a more distant *Kayfunavirus* relatives and other small *Siphoviridae* species (39). BLAST database hits with high percent identity at the nucleotide level compared to CROW1, CROW2, CROW3, and CROW4 revealed phage EcSW4, a related *Kayfunavirus* isolated from polluted stream water in Brazil with its terminal repeats identified (40). Genomes were compared (Figure 3B), and the tail spikes were at least 98% identical to that of EcSW4. Although no significant difference was observed in the tail spikes, notable differences were present in an ATP-dependent DNA ligase, gp5.5-like host HNS inhibition gene product, and internal virion protein. Additionally, CROW1 contained an extra HNH endonuclease protein that was not present in any of these closely related phages.

**Figure 3.**
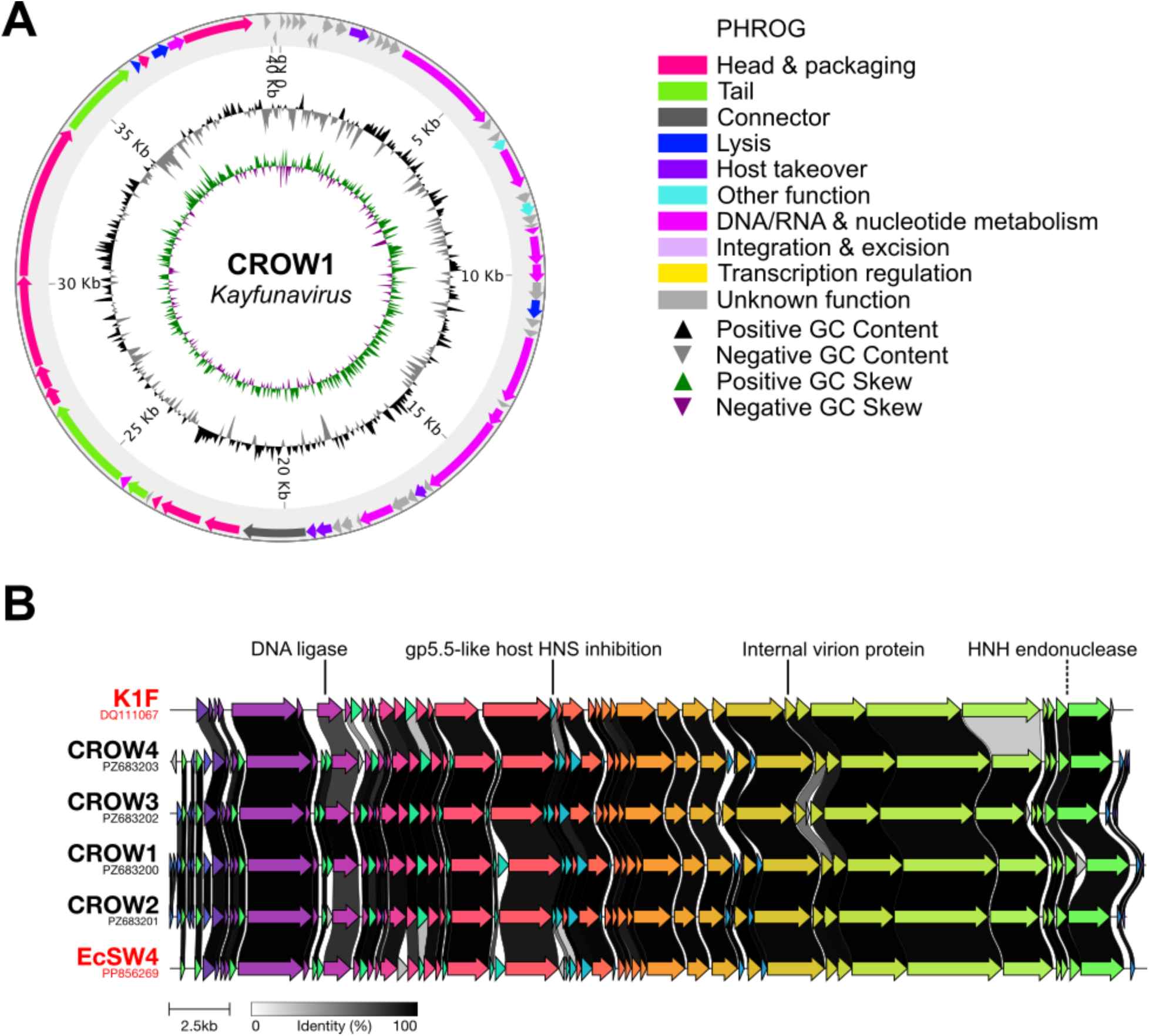
Genomic landscape and coding region comparison of *Kayfunavirus* bacteriophages. (**A**) The annotated genome of CROW1 generated by phold. Coding regions are categorized into color-coded groups according to the PHROG (Prokaryotic Virus Remote Homologous Groups) library. Positive/negative GC content and skew are shown on a radial scale, and genome locations are labeled at 5 kilobase (Kb) intervals. (**B**) Comparison of proteins between discovered (black) and reference (red) bacteriophage genomes. Amino acid percent identity (grayscale) is displayed as a linker between gene products. The lowest identity and extra/absent gene products are labeled with a solid and dashed line respectively.

### Genomic analysis of Vectrevirus bacteriophages

The only *Vectrevirus* bacteriophage that was isolated, CROW5, has a genome size of 45,117 base pairs consisting of 45.08% GC content and no significant stretches of GC skew changes. All 61 ORFs are oriented in the same direction, and 26 of them were annotated with an unknown function (Figure 4A). Structural, metabolic, and lysis genes were present, but tRNA genes were absent. CROW5 was directly compared to K1-5, a closely related bacteriophage that has two ORFs in the tail fiber protein region that allows for infection of both K1 and K5 capsule types of *E. coli* (41). In addition, another *Vectrevirus*, K1E, only possesses one of these ORFs corresponding to the K1 endosialidase tail fibers and can therefore only infect that capsule type (42). This gene product was strikingly similar between CROW5, K1-5, and K1E, however CROW5 has different and slightly shorter K5 lyase tail fiber gene than K1-5 (Figure 4B). CROW5 is >95% identical to K1-5, both of which share similarity to SP6, a Salmonella phage, implying each of these has diverged from T7 phages (43,44). Besides the tail fibers, low similarity scores were observed for the structural major head protein and head scaffolding protein. Differences in amino acid identity were also present between multiple acetyltransferases, a nucleotide kinase and an ATP-dependent DNA ligase. Additionally, two extra HNH endonuclease gene products were present in CROW5 that were not seen in K1-5.

**Figure 4.**
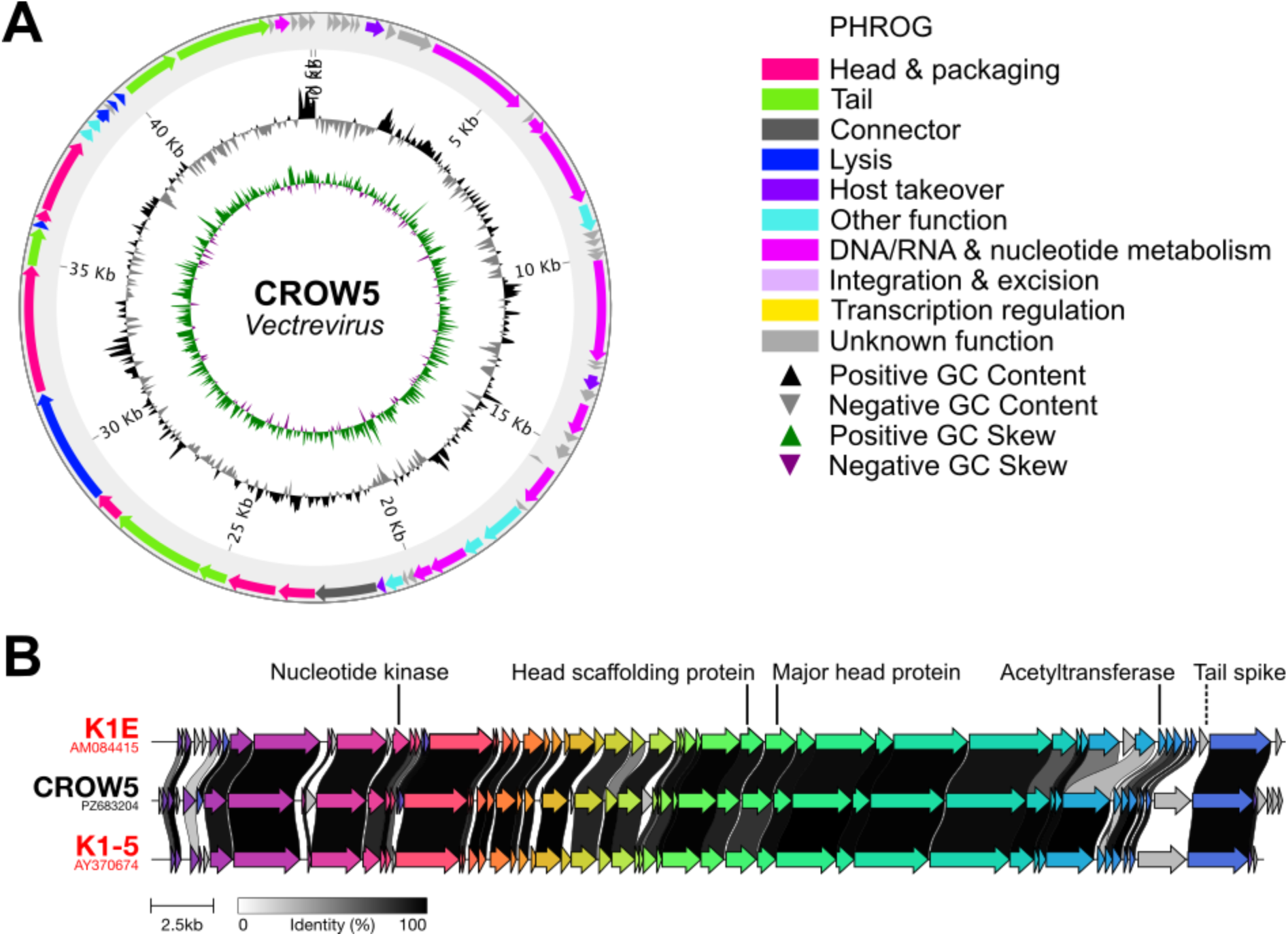
Genomic landscape and coding region comparison of *Vectrevirus* bacteriophages. (**A**) The annotated genome of CROW5 generated by phold. Coding regions are categorized into color-coded groups according to the PHROG (Prokaryotic Virus Remote Homologous Groups) library. Positive/negative GC content and skew are shown on a radial scale, and genome locations are labeled at 5 kilobase (Kb) intervals. (**B**) Comparison of proteins between discovered (black) and reference (red) bacteriophage genomes. Amino acid percent identity (grayscale) is displayed as a linker between gene products. The lowest identity and extra/absent gene products are labeled with a solid and dashed line respectively.

### Genomic analysis of Kuravirus bacteriophages

Both bacteriophages belonging to the genus *Kuravirus*, CROW6 and CROW7, had very similar sized genomes of 77,425 and 77,474 base pairs. Using CROW6 as a representative genome, the GC content is 42.18% with a change in GC skew that corresponds with the direction of two major groups of ORFs. A total of 152 ORFs are present, however only 53 were able to be annotated and the majority of hypothetical ORFs clustered into the same region of the genome (Figure 5A). Head and tail genes were identified, several genes related to DNA/RNA metabolism and amino-acid modifying enzymes were also present, along with a single tRNA gene. Although these bacteriophages were independently isolated from sludge and influent, their sequences are 99.9% identical to one another. One close relative based on ViPTree, CHD5UKE1, was isolated from wastewater in Braunschweig, Germany using a clinically derived uropathogenic *E. coli* strain (45). Another top hit on the proteomic tree was phiEco2, a bacteriophage isolated in Tbilisi, Georgia that appears to share several genes that are also present in a *Pseudomonas aeruginosa* phage (46). Some unique start and stop codon usage was also reported, along with the sequence of terminal repeats that aided in the orientation of CROW6 and CROW7 genomes. Bacteriophages CROW6 and CROW7 were more closely related to SU10 than phiEco32, with an in-depth comparison already conducted between the two. Structural head proteins were very well conserved between each of these *Kuravirus* species. The elongated C3 head morphotype is not a product of the major capsid gene length, however unique features of the head scaffolding proteins could aid in the formation of this structure (47). The more divergent tail fiber proteins play a major role in the binding of host receptors and induce conformational changes in the nozzle and short tail fibers that are required to penetrate the host membrane (48). Unique tail fiber proteins are not present in CROW6 and CROW7 (>99% similarity to CHD5UKE1), but other differences have the potential to impact the cascade of interactions affecting host attachment and takeover that could alter host specificity (Figure 5B). These notable differences based on low similarity scores included a polymerase, anti-restriction nuclease, D5-like transcriptional regulator, membrane protein, and ribosomal protein S6 glutaminyl transferase. Another clear difference is the absence of a phosphatase and HNH endonuclease in CROW6 and CROW7 that was present in the reference genomes.

**Figure 5.**
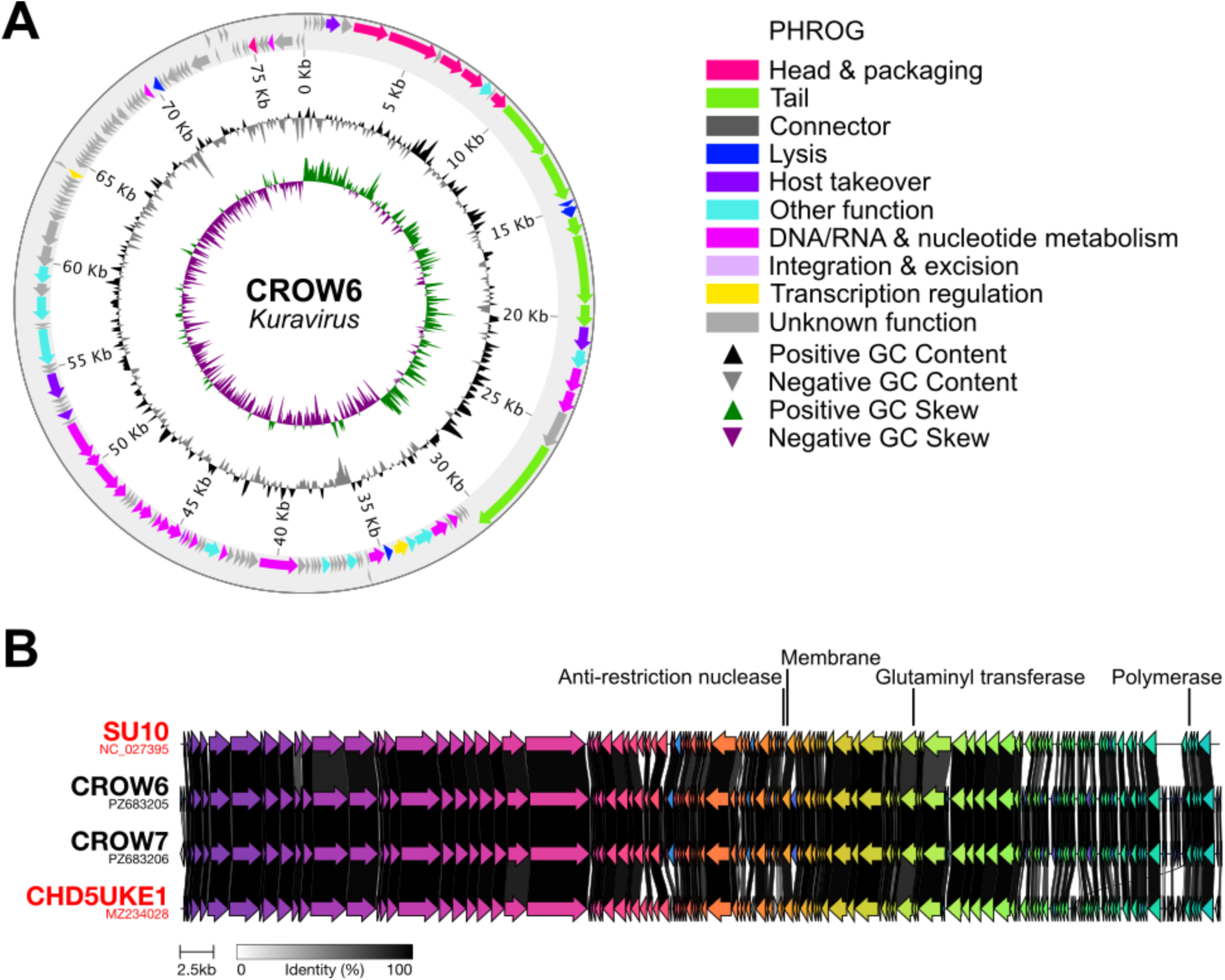
Genomic landscape and coding region comparison of *Kuravirus* bacteriophages. (**A**) The annotated genome of CROW6 generated by phold. Coding regions are categorized into color-coded groups according to the PHROG (Prokaryotic Virus Remote Homologous Groups) library. Positive/negative GC content and skew are shown on a radial scale, and genome locations are labeled at 5 kilobase (Kb) intervals. (**B**) Comparison of proteins between discovered (black) and reference (red) bacteriophage genomes. Amino acid percent identity (grayscale) is displayed as a linker between gene products. The lowest identity gene products are labeled with a solid line.

### Genomic analysis of Sugarlandvirus bacteriophages

All three bacteriophages isolated with the *Klebsiella pneumoniae* host were classified into the genus *Sugarlandvirus*. CROW8, CROW9, and CROW10 had the largest genomes (111,559, 111,600, or 111,359 base pairs respectively) compared to the bacteriophages isolated on the *E. coli* strain. Using CROW10 as an example, 185 ORFs were identified with 120 of them lacking annotation calls (Figure 6A). Most ORFs with an unknown function appeared to be very short and clustered together. Genes with structural functions such as the head, tail, and connector genes were present, along with several genes related to DNA/RNA metabolism, lysis, and host takeover. In addition, 25 tRNA genes were localized in a single cluster within the genome. CROW8, CROW9, and CROW10 had very similar genomes based on a 92.1-96.0% nucleotide identity to one another. These are not a unique species, showing significant homology (>95% nucleotide identity) to several related *Sugarlandvirus* that have been characterized previously. A closely related *Sugarlandvirus* is phage IME260, which was isolated from sewage collected from a hospital in China and infects several antibiotic-resistant *K. pneumoniae* strains (49). The genomic location of IME260 terminal repeats aided us in orienting the genomes of CROW8, CROW9, and CROW10. Several other phages with high similarity at the nucleotide level are present in the NCBI database such as Kpl_K32PH164C1 and vB_Kpn_K30lambda2.2, two *Sugarlandvirus* phages that were isolated from sewage samples in Spain (50). These bacteriophages possess divergent receptor binding proteins that allow for a broad capsule-dependent host range despite lacking depolymerase domains. The receptor binding proteins of CROW8, CROW9, and CROW10 were at least 94% identical to each of these two other phages supporting their broad host-range potential. Further comparisons revealed a myriad of differences between hypothetical proteins and a few structural proteins (Figure 6B). Three gene products corresponding to tail formation varied between strains: the tail fiber, straight tail fiber, and tail length tape measure protein. Another structural protein containing an immunoglobulin-like domain involved in packaging also showed some divergence from these other closely related bacteriophages.

**Figure 6.**
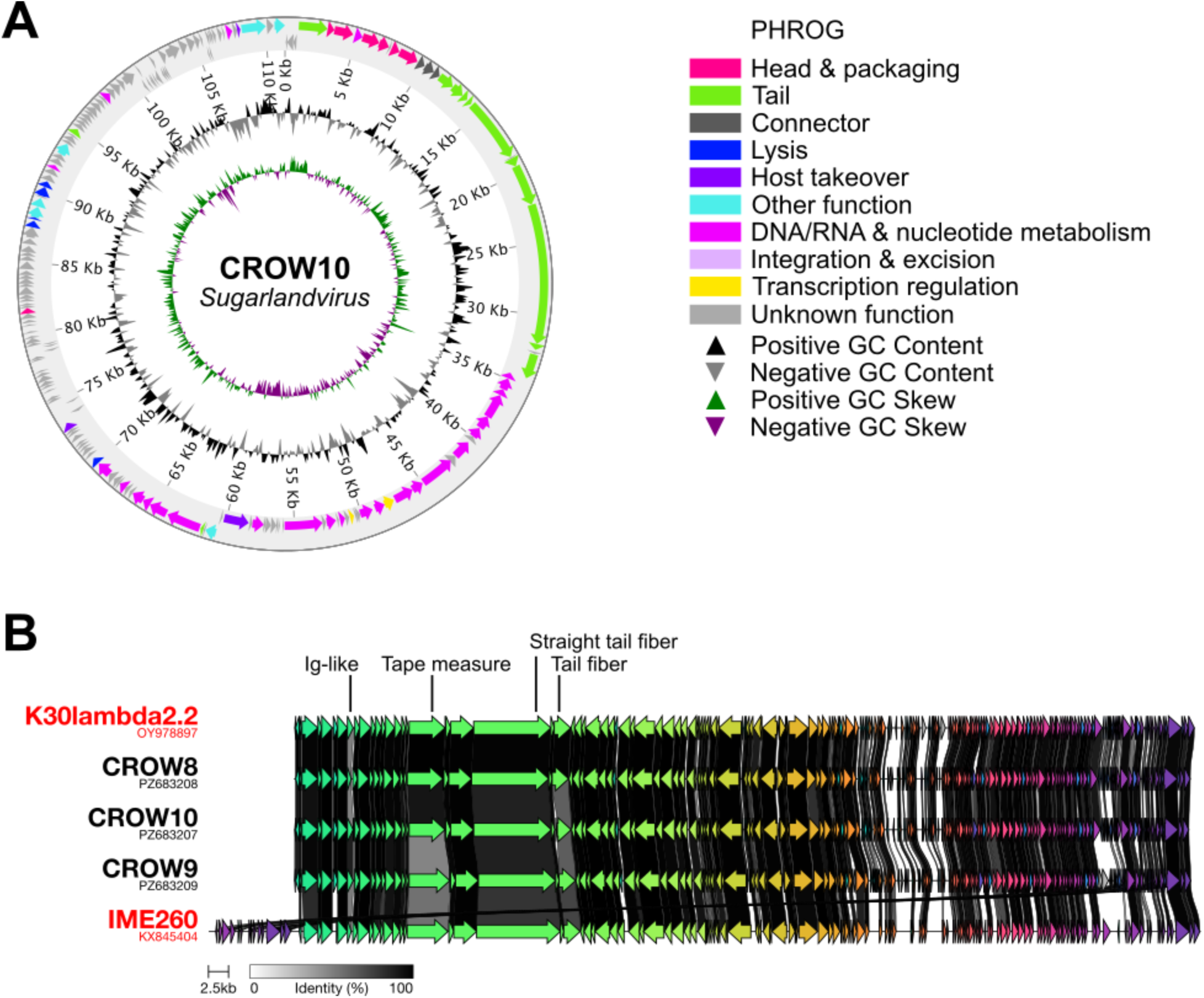
Genomic landscape and coding region comparison of *Sugarlandvirus* bacteriophages. (**A**) The annotated genome of CROW10 generated by phold. Coding regions are categorized into color-coded groups according to the PHROG (Prokaryotic Virus Remote Homologous Groups) library. Positive/negative GC content and skew are shown on a radial scale, and genome locations are labeled at 5 kilobase (Kb) intervals. (**B**) Comparison of proteins between discovered (black) and reference (red) bacteriophage genomes. Amino acid percent identity (grayscale) is displayed as a linker between gene products. The lowest identity gene products are labeled with a solid line.

## 6. Discussion

### Wastewater from Ottawa is an abundant source of Escherichia coli & Klebsiella pneumoniae bacteriophages

Wastewater taken from the water resource recovery facility and university sewer shed in Ottawa during the Summer and Fall of 2025 was screened for the presence of bacteriophages against hypervirulent strains of *E. coli* O1:K1:H7 and *K. pneumoniae* K3. Ten bacteriophages were isolated and characterized using electron microscopy, long-read sequencing, taxonomic classification, and genome annotation. Bacteriophages with *Acinetobacter baumannii* and *Pseudomonas aeruginosa* hosts have been isolated from Ottawa wastewater previously, however only from influent using enrichment techniques (51,52). Many phages are isolated from diluted water sources, but some examples using sludge exist (53–55). Phages from this study were seemingly abundant in all sample types (i.e. sewer shed wastewater, influent wastewater, and primary sludge) and were easily isolated using classical centrifugation and filtration techniques. Isolation did not require enrichment and based on the high frequency of samples with phages large panels of hosts could be screened using these samples (Supplemental Figure 1). Overall, sequencing and morphological analysis revealed an intrasample diversity of bacteriophage, especially considering only two hosts and a short sampling timeframe were employed. That said, intersample diversity of bacteriophages was minimal, with all three sample types, despite originating from different steps in the wastewater reclamation process, yielded similar phages. *Sugarlandvirus* phages were found in all three types, and all *E. coli* phages were isolated independently from both sludge and influent except for *Vectrevirus* (found in sludge but not in influent). The lack of sample specificity speaks to the ubiquitous nature of phages across multiple stages of sewer shed, however our sample size does not allow us to explore if each unique environment provides different evolutionary pressures.

### Current and developing applications of wastewater-derived bacteriophages

There are several examples of phage therapy being successfully used to treat *E. coli* and *K. pneumoniae* infections in humans over recent years (56,57). Several *Kayfunavirus* species have been reported to infect a broad range *E. coli* serogroups, including human clinical isolates of O45 and O25, and avian pathogenic *E. coli* O1/O2 strains (58–60). A wide range of water sources continue to be a rich source of phages for this genus and should continue to be explored in an effort to combat the global concern of antimicrobial resistance (61).

Very few *Vectrevirus* phages have been isolated and characterized to date. Several were isolated from wastewater in a study conducted in Turkey where they were shown to infect a clinical multidrug-resistant strain of *E. coli*, and another study in Germany where they were screened against a panel of *E. coli* strains (45,62). Some species previously classified as *Vectrevirus* have been reassigned to *Rodentiumvirus* (63), potentially due to unique characteristics that allow these viruses to infect *Citrobacter rodentium*, a model organism for enteropathogenic *E. coli* (64).

Multiple other examples of wastewater being a source of *Kuravirus* exist, including one from activated sludge, further supporting the feasibility of this less common sample type when screening for phage (65). *Kuravirus* bacteriophages have also been shown to infect multiple avian pathogenic *E. coli* strains and be effective against their biofilms (66). Similar phages reduced the concentration of viable Shiga toxin-producing E. coli in milk models highlighting their potential in agricultural and food safety applications (67). Bacteriophages belonging to the *Kuravirus* genus are growing in promise and recognition; noted by the reclassification of some viruses to *Kuravirus* based on 12 core proteins (63). Furthermore, they have shown recent success in a phage therapy cocktail used in a human clinical trial against urinary tract infections (68).

*Sugarlandvirus* phages have a relatively broad host range for *K. pneumoniae* strains and can infect a range of multidrug-resistant strains with various capsule types (69,70). Despite the large range of hosts that are susceptible to these viruses, and general promise for *K. pneumoniae* phages in human clinical trials, there have been none using a *Sugarlandvirus* yet (71). *Sugarlandvirus* has been shown to effectively lower bovine *K. pneumoniae* burden in mouse models and reduce contamination on vegetable and poultry products (72,73), suggesting it has promise as a therapeutic phage.

### Phage biobanks are needed to expedite bacteriophage research

Safety is a primary concern when considering FDA approval for bacteriophages in clinical trials, and widespread efficacy is a large speedbump for interested pharmaceutical parties. Despite these challenges, initial results are promising enough that effort is being directed into building frameworks for faster approval that go hand in hand with the resurgence of bacteriophage isolation and characterization by scientists globally (74). As phage therapy workflows continue to develop, a need for standardized phage biobanking has become apparent (75). Due to the nature of host specificity, screens are usually conducted to find a phage that is effective against any given clinical bacterial isolate. Having access to a panel of well characterized phages streamlines the testing required to go from diagnosis to treatment (76). This work has generated long-term cryogenic stocks of ten purified bacteriophages and contributed to the database of sequences in GenBank. These phages were isolated directly from small-volume samples, supporting the use of centralized municipal wastewater as a diverse and accessible source of bacteriophages for clinical and agricultural research, rather than relying on facility-specific sites or fully developed centralized biobanks.

## 7.3 Acknowledgements

The authors acknowledge the Electron Microscopy Core (RRID: SCR_025398) funded by the University of Ottawa, Brain-Heart Interconnectome (BHI) via Canadian First Research Excellence Fund (CFREF). Special thanks to Kieran Furlong at the University of Ottawa for discussions about bacteriophage sequencing and purification.

