## Supplemental Figures 1 & 2 for "Municipal Wastewater in Ottawa, Canada contains a reservoir of bacteriophages with clinically relevant *Escherichia coli* and *Klebsiella pneumoniae* hosts"

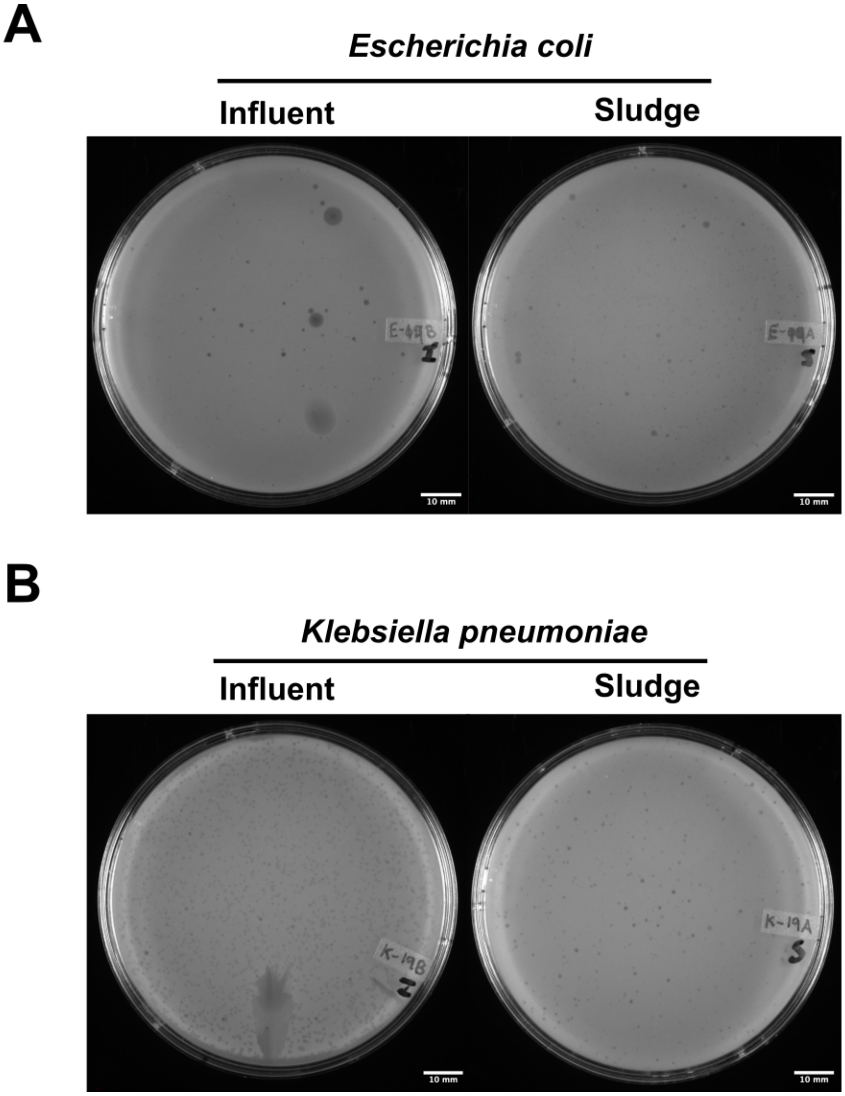


**Supplemental Figure 1.** Direct isolation of bacteriophage plaques on *Escherichia coli* and *Klebsiella pneumoniae* hosts.

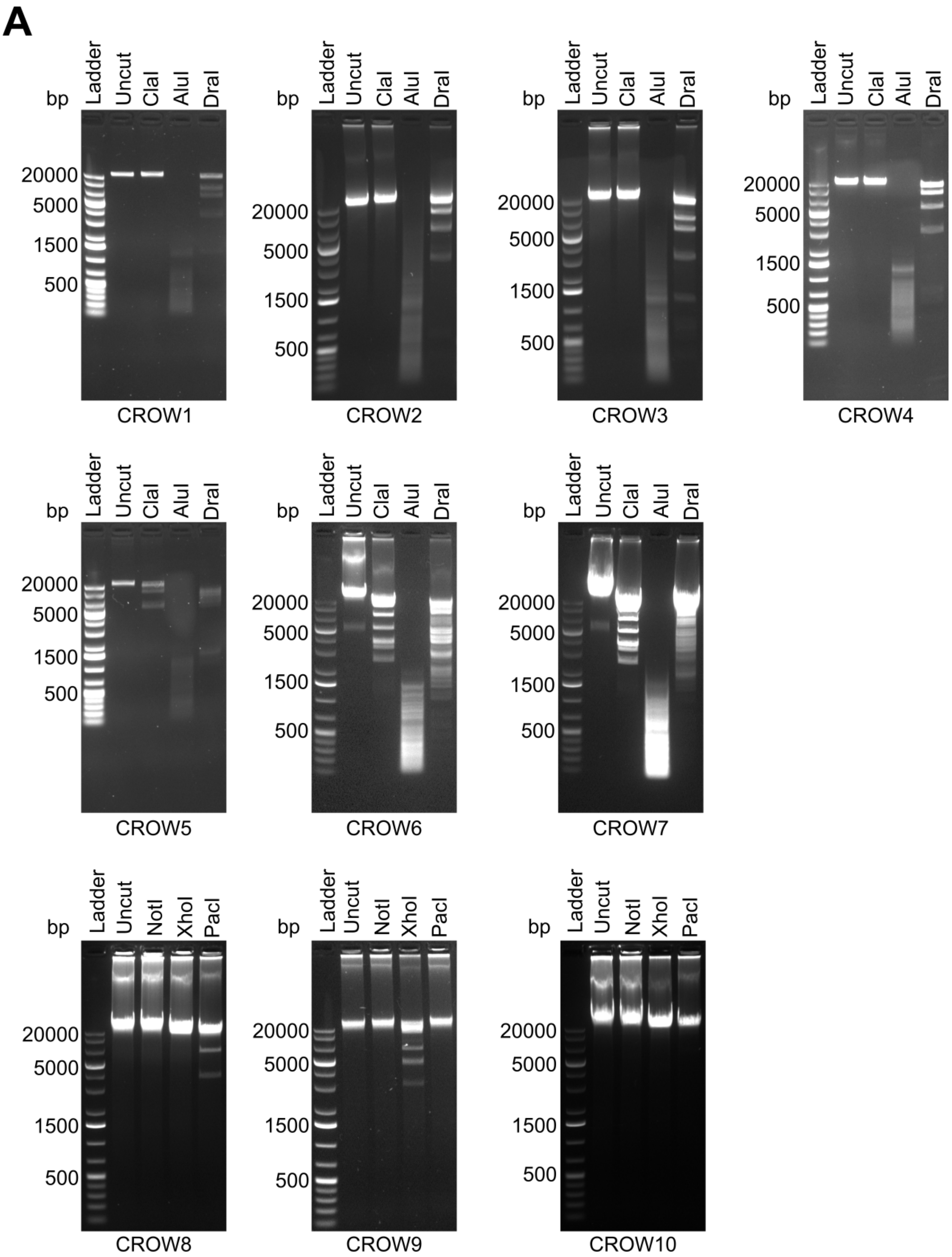


**Supplemental Figure 2.** Restriction Fragment Length Polymorphism Analysis.
